# SARS-CoV-2 manipulates the SR-B1-mediated HDL uptake pathway for its entry

**DOI:** 10.1101/2020.08.13.248872

**Authors:** Congwen Wei, Luming Wan, Qiulin Yan, Xiaolin Wang, Jun Zhang, Yanhong Zhang, Jin Sun, Xiaopan Yang, Jing Gong, Chen Fan, Xiaoli Yang, Yufei Wang, Xuejun Wang, Jianmin Li, Huan Yang, Huilong Li, Zhe Zhang, Rong Wang, Peng Du, Yulong Zong, Feng Yin, Wanchuan Zhang, Yumeng Peng, Haotian Lin, Rui Zhang, Wei Chen, Qi Gao, Yuan Cao, Hui Zhong

## Abstract

The recently emerged pathogenic severe acute respiratory syndrome coronavirus 2 (SARS-CoV-2) has spread rapidly, leading to a global COVID-19 pandemic. Binding of the viral spike protein (SARS-2-S) to cell surface receptor angiotensin-converting enzyme 2 (ACE2) mediates host cell infection. In the present study, we demonstrate that in addition to ACE2, the S1 subunit of SARS-2-S binds to HDL and that SARS-CoV-2 hijacks the SR-B1-mediated HDL uptake pathway to facilitate its entry. SR-B1 facilitates SARS-CoV-2 entry into permissive cells by augmenting virus attachment. MAb (monoclonal antibody)-mediated blocking of SARS-2-S-HDL binding and SR-B1 antagonists strongly inhibit HDL-enhanced SARS-CoV-2 infection. Notably, SR-B1 is co-expressed with ACE2 in human pulmonary and extrapulmonary tissues. These findings revealed a novel mechanism for SARS-CoV-2 entry and could provide a new target to treat SARS-CoV-2 infection.

## Introduction

Since the outbreak of COVID-19 caused by severe acute respiratory syndrome coronavirus 2 (SARS-CoV-2), over half million people have died and the total number of global confirmed cases was now over 10 million as of July 28, 2020^1^. Coronavirus tropism is predominantly determined by the interaction between coronavirus spikes (S) and their corresponding host receptors^2,3^. The S protein of SARS-CoV-2 (SARS-2-S) can be cleaved into the S1 (SARS-2-S1) and S2 (SARS-2-S2) subunits, which are responsible for receptor recognition and membrane fusion^4^. Binding of SARS-CoV-2 to ACE2 occurs via the C-terminal domain (also called receptor binding domain [RBD]) of SARS-2-S1^5^. The observed incomplete inhibition of ACE2 antibody and the high neutralization potency of mAb (monoclonal antibody) targeting the N-terminal domain (NTD) of SARS-2-S1 suggest that SARS-CoV-2 might may use other (co)receptors, or auxiliary proteins or even other mechanisms for host cell entry^6^.

Scavenger receptor B type 1 (SR-B1) is a cell surface HDL receptor that mediates the selective uptake of HDL-CE (cholesteryl esters) and other lipid components of receptor-bound HDL particles, including free cholesterol (FC), triglycerides (TGs), phospholipids (PLs), α-tocopherol and vitamin E^7^. This cholesterol delivery system is well operated in isolated hepatocytes, fibroblasts, adipocytes, macrophages, adrenal, ovarian and testicular Leydig cells^8^. Interestingly, alveolar type II cells also express SR-B1, which is responsible for vitamin E uptake preferentially from HDL^9,10^. SR-B1 has emerged as a critical receptor that affects HCV entry^11^. However, the potential role of SR-B1 in SARS-CoV-2 infection has yet to be evaluated.

## Results

Due to the importance of SARS-2-S in mediating the infection of target cells, a search for cholesterol-regulated motifs in the primary sequence of SARS-2-S was conducted and 6 putative cholesterol recognition amino acid consensus (CRAC) motifs adjacent to the inverted cholesterol recognition motif called CARC were identified^12^ (Fig. 1a). We then evaluated the ability of SARS-2-S to associate with cholesterol. The purity of SARS-2-S, SARS-2-S1 and SARS-2-S2 protein was validated by SDS-PAGE (Extended Data Fig. 1a). MST assay showed that SARS-2-S could bind to cholesterol with an EC_50_ of 187.6±120.5 nM (Fig. 1b). Specifically, SARS-2-S did not exhibit any binding with campesterol and epicholesterol, two structurally distinct sterols with additional methyl and hydroxyl groups (Fig. 1c). In a competition assay, unlabeled SARS-2-S protein had an IC_50_ of 821±69.8 nM for cholesterol (Extended Data Fig. 1b). In subsequent analyses, we evaluated the interaction of SARS-2-S, SARS-2-S1 and SARS-2-S2 with HDL particles. While SARS-2-S and SARS-2-S1 bound to HDL (Fig. 1d), SARS-2-S2 failed to bind HDL (Fig. 1e). Furthermore, immunoprecipitation analysis revealed that SARS-2-S1 did not bind to ApoA1 (Extended Data Fig. 1c). Taken together, these results strongly support the notion that the SARS-2-S has a specific affinity for cholesterol and HDL. We then attempted to dissect the interacting determinants in SARS-2-S through an in vitro binding assay. Among the four cholesterol binding peptides encompassing the CARC-CRAC region, three putative cholesterol recognition regions were observed to reside in the CTD of SARS-2-S1, while one motif was located in the RBD (Fig. 1f). In consistence with the MST data, the 2 cholesterol recognition motifs in SARS-2-S2 did not show any binding affinity with cholesterol (Fig. 1f).

**Fig. 1.**
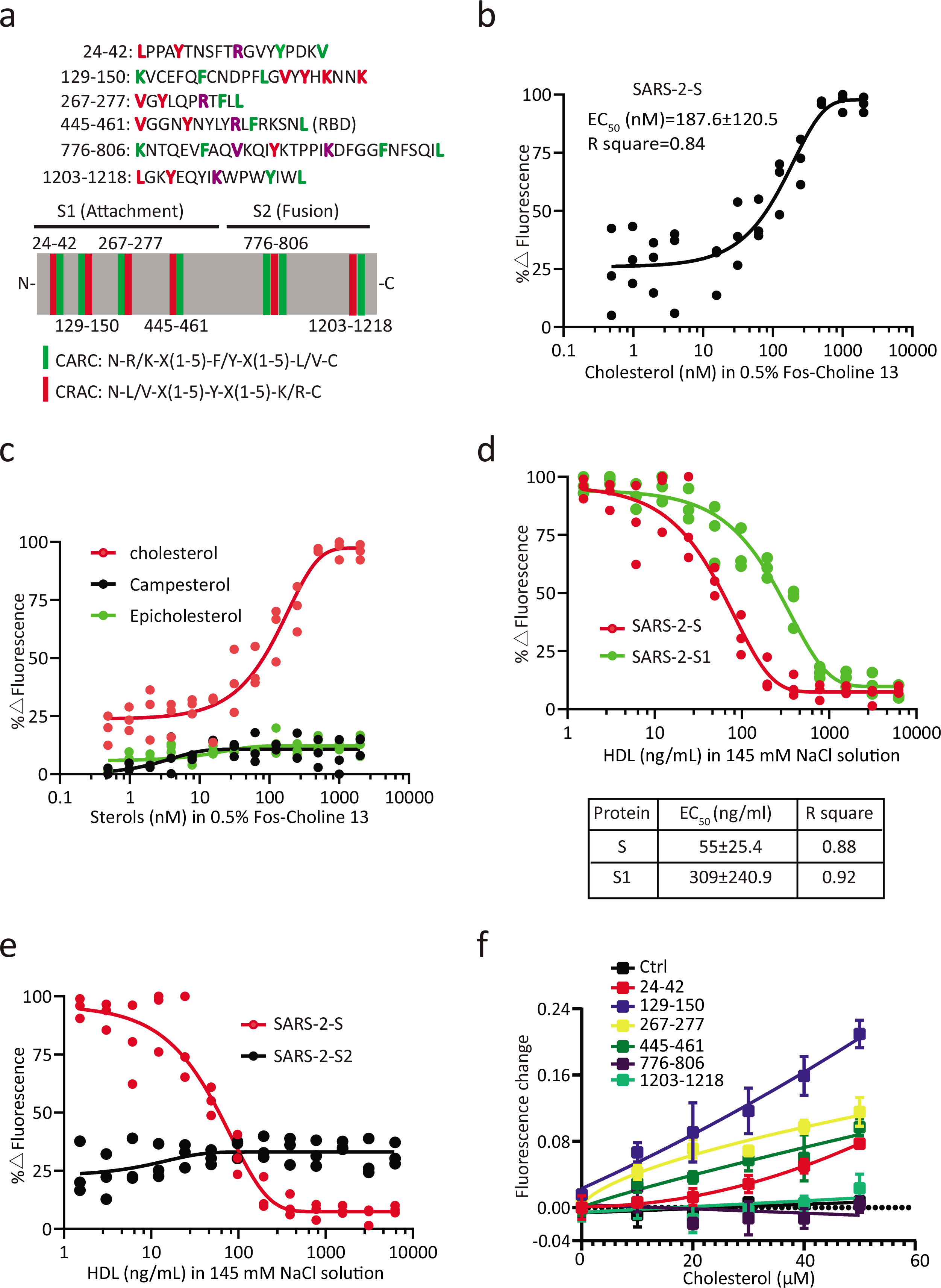
The spike protein of SARS-CoV-2 binds to cholesterol and HDL. a, Schematic illustration of the cholesterol-binding motif in SARS-2-S. Crucial amino acid residues in the CARC motif are highlighted in red, crucial amino acid residues in the CRAC motif are highlighted in green, and shared amino acid residues are highlighted in purple. The green stick indicates the CARC motif, and the red stick indicates the CRAC motif. b,c, MST analysis of interactions between SARS-2-S and cholesterol (b) or campesterol and epicholesterol (c) (n= 3). Data were derived from the effect of cholesterol on fluorescence decay of fluorescently labeled SARS-2-S. EC_50_ was determined by Hill slope. d,e, MST analysis of interactions between SARS-2-S, SARS-2-S1 (d) or SARS-2-S2 (e) with HDL (n= 3). Data were derived from the effect of HDL on fluorescence decay of fluorescently labeled proteins as evaluated with MST. EC_50_ was determined by Hill slope. f, Titration curves displaying changes in the intrinsic fluorescence of the CARC-CRAC peptide (5 μM) in the presence of increasing concentrations of cholesterol (n=3). Data are mean ± SEM.

We then want to determine whether the association of SARS-2-S1 with HDL affects SARS-CoV-2 entry. To this end, the capacity of HDL in affecting the entry of SARS-2-S-pseudovirus (SARS-CoV-2pp) was explored (Fig. 2a). HDL significantly enhanced SARS-CoV-2pp entry in a dose dependent manner (Fig. 2b), and similar results were observed in Huh-7 cells infected with authentic SARS-CoV-2 (Fig. 2c). To investigate the role of HDL in SARS-2-S cell surface attachment, cells were incubated with SARS-2-S in solutions containing HDL at RT for 30 min and immunolabeled for SARS-2-S after washing. Quantitative analysis of the expression rate (Fig. 2d) and mean fluorescent intensity (MFI; Fig. 2e) revealed that HDL induced a substantial increase in cell surface SARS-2-S attachment in a dose dependent manner.

**Fig. 2.**
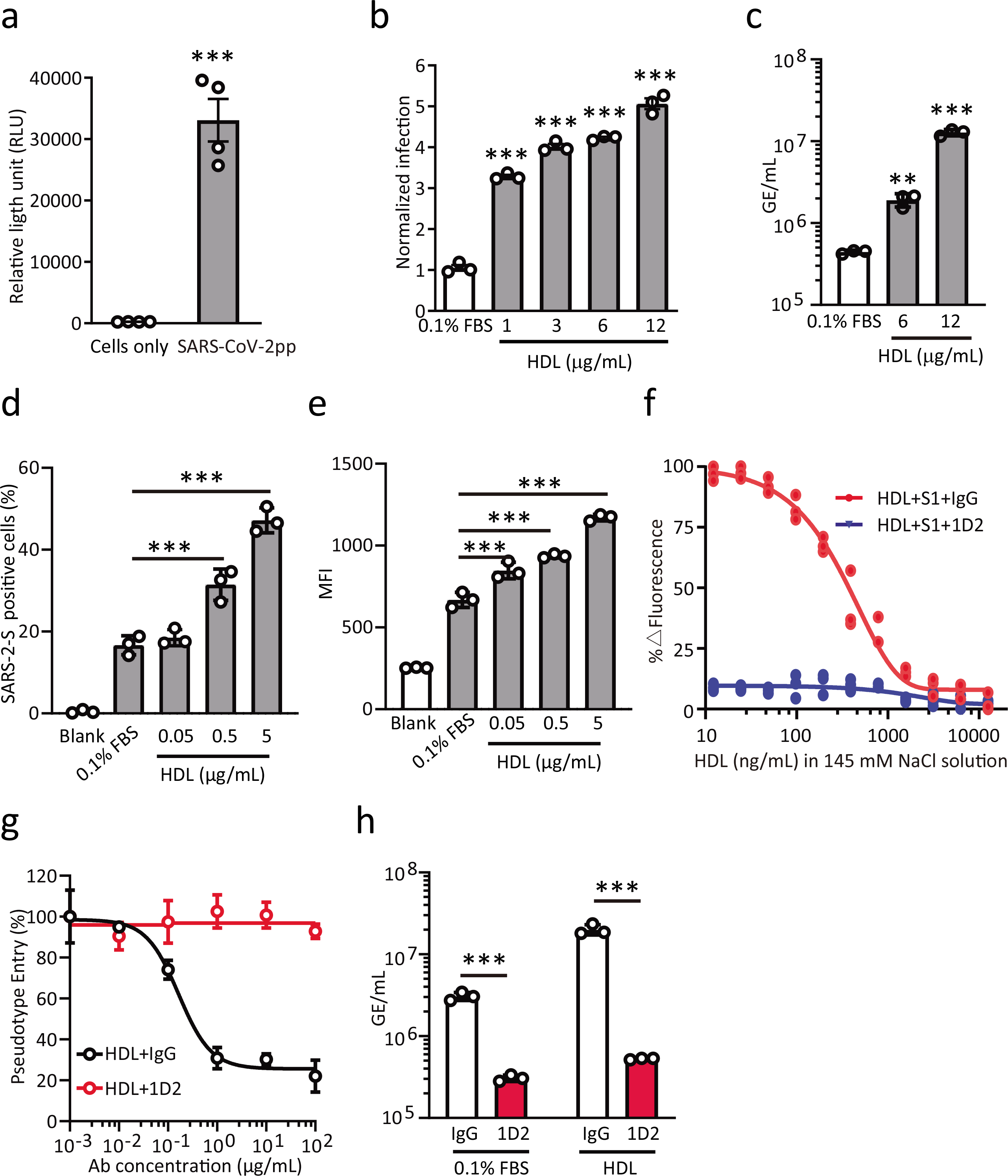
Binding of SARS-2-S-HDL enhances SARS-CoV-2 entry. (a) Huh-7 cells were inoculated with SARS-2-Spp in 10% FBS and pseudotyped viral entry was analyzed by luciferase activity at 48 h after infection (n=3). Data are mean ± SEM. ***P < 0.001 by Two-tailed Student’s t-test. b, Huh-7 cells were inoculated with SARS-CoV-2pp in solutions containing HDL at the indicated concentrations and pseudotyped viral entry was analyzed by luciferase activity at 48 h after infection (n=3). Signals obtained in 0.1% FBS were used for normalization. Data are mean ± SEM. ***P < 0.001 by one-way ANOVA and Bonferroni’s post hoc analysis. c, Huh-7 cells were infected with SARS-CoV-2 (MOI = 0.001) in solutions containing HDL at the indicated concentrations for 1 h and the cells were harvested to detect the gene copy number of virus with qRT-PCR at 24 h after infection (n=3). Data are mean ± SEM. **P < 0.01, ***P < 0.001 by one-way ANOVA and Bonferroni’s post hoc analysis. d,e, HeLa-hACE2 cells were exposed to SARS-2-S in solutions containing HDL at the indicated concentrations at RT. After 30 min, the cells were washed and detached before immunolabeling for flow cytometry (n=3). Percentage of SARS-2-S positive cells (d) and mean fluorescence intensity (MFI) of SARS-2-S (e) on the cell surface was quantified with isotype and SARS-2-S staining. Data are mean ± SEM. ***P < 0.001 by one-way ANOVA and Bonferroni’s post hoc analysis. f, Preincubation of 1D2 (200 nM) with SARS-2-S1 (100 nM) and the effect of HDL on fluorescence decay of fluorescently labeled SARS-2-S1 was evaluated with MST (n=3). EC_50_ was determined by Hill slope. g, Huh-7 cells preincubated with serially diluted 1D2 were inoculated with SARS-2-Spp in solutions containing 6 μg/mL HDL and pseudotyped viral entry was analyzed by luciferase activity at 48 h after infection (n=3). Signals obtained without 1D2 were used for normalization. Data are mean ± SEM. h, Huh-7 cells were infected with SARS-CoV-2 (MOI = 0.001) preincubated with 1D2 (167 nM) in solutions containing 6 μg/mL HDL at the indicated concentrations for 1 h and the cells were harvested to detect the gene copy number of virus with qRT-PCR at 24 h after infection (n=3). Data are mean ± SEM. ***P < 0.001 by two-tailed Student’s t-tests.

In a previous study, the mAb 1D2 was isolated from convalescent COVID-19 patients and shown to exhibit high neutralization potency against SARS-CoV-2^6^. Lys147, Lys150 and Tyr145, which are located in the HDL-binding motif of SARS-2-S, were identified as important antigenic sites for 1D2^6^. We then examined the ability of this mAb to interfere with HDL binding to SARS-2-S1. Preincubation of 1D2 with SARS-2-S1 before the addition of HDL completely blocked SARS-2-S1-HDL binding (Fig. 2f), strongly reduced HDL-enhanced SARS-CoV-2pp entry (Fig. 2g) and authentic SARS-CoV-2 infection (Fig. 2h). Collectively, these data show that HDL enhances SARS-CoV-2 infectivity by interacting with the NTD of SARS-2-S1.

As SR-B1 is a cell-surface receptor that binds HDL and plays a central role in HDL endocytosis and cholesterol efflux, we wondered whether the HDL-mediated enhancement of SARS-CoV-2 infectivity occurs through SR-B1. To test this possibility, infection assays were performed using two SARS-CoV-2pp permissive cell lines, Huh-7 and Vero E6. Huh-7 cells displayed higher surface (Extended Data Fig.2a) and intracellular (Extended Data Fig.2b) SR-B1 and ACE2 protein levels than Vero cells. Notably, ectopic SR-B1 overexpression specifically enhanced Huh-7 susceptibility to SARS-CoV-2pp, but not to unrelated VSV-Gpp (Fig. 3a and Extended Data Fig.2c). In contrast, SR-B1 downregulation significantly reduced HDL-mediated enhancement of infection (Fig. 3b and Extended Data Fig. 2d). We next sought to determine whether SR-B1 overexpression renders cells more susceptible to infection with authentic SARS-CoV-2. The RNA load of SARS-CoV-2 in Huh-7 cells transfected with siSR-B1 (Extended Data Fig. 2e) was approximately 10 times lower than that observed in control cells (Fig. 3c), while SR-B1 overexpression (Extended Data Fig. 2f) in Vero E6 cells potently increased SARS-CoV-2 infection (Fig. 3d).

**Fig. 3.**
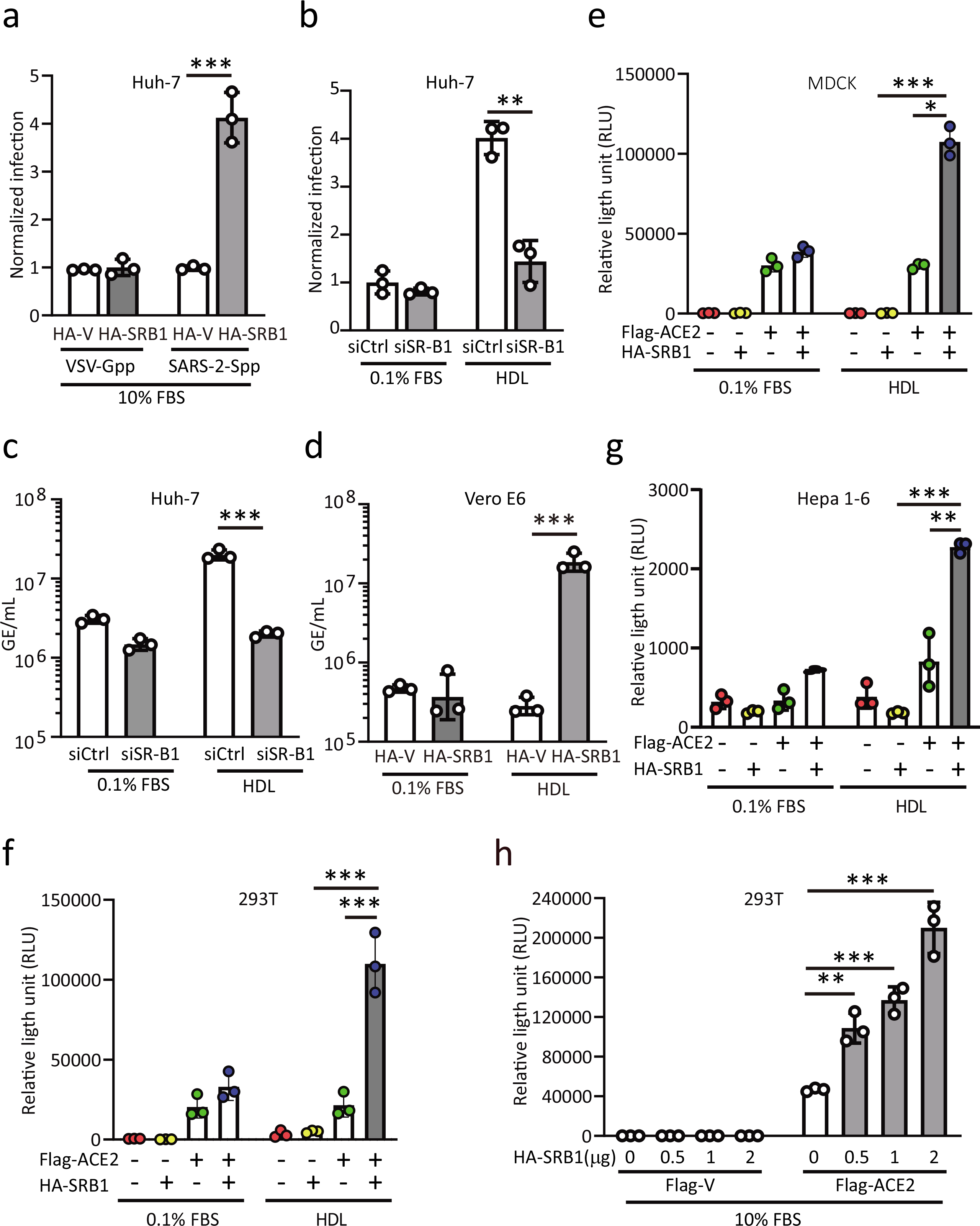
SR-B1 expression confers susceptibility to SARS-CoV-2 infection. a, Huh-7 cells transfected with HA-Vector or HA-SR-B1 were inoculated with SARS-2-Spp or VSV-Gpp in 10% FBS and pseudotyped viral entry was analyzed by luciferase activity at 48 h after infection (n=3). Signals obtained in cells transfected with HA-Vector were used for normalization. Data are mean ± SEM. ***P < 0.001 by two-tailed Student’s t-tests. b, Control (siCtrl) or SR-B1 knockdown (siSR-B1) Huh-7 cells were inoculated with SARS-CoV-2pp in solutions containing 0.1% FBS or 6 μg/mL HDL and pseudotyped viral entry was analyzed by luciferase activity at 48 h after infection (n=3). Signals obtained in siCtrl cells with 0.1% FBS were used for normalization. Data are mean ± SEM. **P < 0.01 by two-tailed Student’s t-tests. c,d, Huh-7 cells transfected with siSR-B1 (c) or Vero E6 cells transfected with a plasmid encoding SR-B1 (d) were infected with SARS-CoV-2 (MOI = 0.001) in solutions containing 0.1% FBS or 6 μg/mL HDL for 1 h. Cells were harvested to detect the gene copy number of virus with qRT-PCR after 24 h (n=3). Data are mean ± SEM. ***P < 0.001 by one-way ANOVA and Bonferroni’s post hoc analysis. e,f,g, MDCK (e), 293T (f) or Hepa 1-6 (g) cells were either transfected with plasmids encoding ACE2, SR-B1 or both and then challenged with SARS-CoV-2pp in solutions containing 0.1% FBS or 6 μg/mL HDL (n=3). Data are mean ± SEM. *P < 0.05, **P < 0.01, ***P < 0.001 by one-way ANOVA and Bonferroni’s post hoc analysis. h, 293T cells transfected with plasmids encoding SR-B1 together with Flag-Vector or Flag-ACE2 were inoculated with SARS-2-Spp in 10% FBS and pseudotyped viral entry was analyzed by luciferase activity at 48 h after infection (n=3). Data are mean ± SEM. **P < 0.01, ***P < 0.001 by one-way ANOVA and Bonferroni’s post hoc analysis.

To further survey the SARS-CoV-2pp susceptibility, MDCK, 293T and Hepa 1-6 cells with very low SARS-CoV-2pp susceptibility and nearly undetectable SR-B1 and ACE2 expression were evaluated (Extended Data Fig. 2a-b). We challenged these cells with SARS-CoV-2pp after being transfected with cDNA vectors encoding human ACE2 and/ or SR-B1 (Extended Data Fig. 2g). MDCK (Fig. 3e) and 293T (Fig. 3f) became highly susceptible to SARS-CoV-2pp upon ACE2 expression, while Hepa 1-6 remained less permissive (Fig. 3g). Although SR-B1 introduction did not render these cells more permissive, the combined SR-B1 and ACE2 expression resulted in higher susceptibility than that observed with ACE2 or SR-B1 expression alone (Fig. 3e, f, g). In particular, SARS-CoV-2pp infectivity was correlated with only SR-B1 expression levels when ACE2 was overexpressed (Fig. 3h and Extended Data Fig. 2h). Of note, the expression of SR-B1 with or without ACE2 did not modify the cell surface expression levels of either of these entry factors at 24 h after transfection (Extended Data Fig. 2i). Hence, our results indicated that SR-B1 is an entry coreceptor of SARS-CoV-2.

Further analysis was performed to assess the colocalization of SARS-2-Spp with SR-B1 via confocal fluorescence microscopy. SARS-CoV-2pp in solutions containing HDL was incubated with cells on ice for 1 min and subsequently visualized their binding to GFP-SR-B1 expressed on the cell surface. Immunostaining results revealed co-localization of SR-B1 with SARS-2-Spp (Fig. 4a). In particular, GFP-SR-B1 positive cells showed increased intensity of intracellular SARS-CoV-2pp staining as compared to GFP-negative cells when we switched the incubation temperature from 4 °C to 37 °C to allow SARS-CoV-2pp entry (Fig. 4b). In agreement with the above results, ectopic SR-B1 expression also led to increased SARS-2-S cell-surface binding stimulated by HDL (Fig. 4c, d). Because SARS-2-S cannot bind to SR-B1 directly (Fig. 4e), our data indicated that SARS-CoV-2 interacts with SR-B1 bridged by HDL during virus entry.

**Fig. 4.**
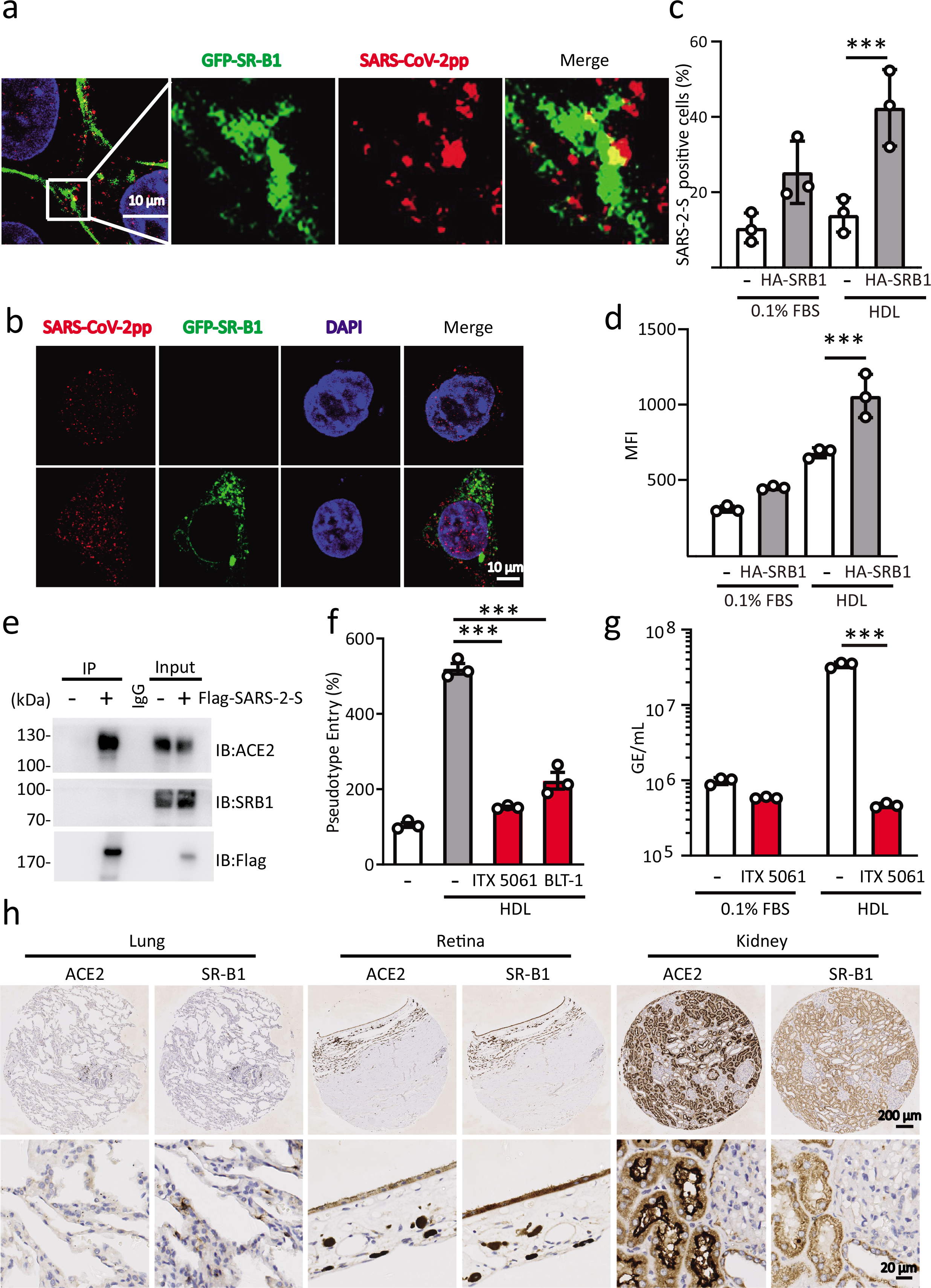
SR-B1 mediates SARS-CoV-2 attachment and entry. a, 293T cells transfected with GFP-SR-B1 were exposed to SARS-CoV-2pp in solutions containing 6 μg/mL HDL at RT for 5 min to attach to the cell surface. All images were obtained by confocal microscopy using a Nikon A1. The scale bar indicates 10 μm. The data shown are representative of two independent experiments. b, 293T cells transfected with GFP-SR-B1 were exposed to SARS-CoV-2pp in solutions containing 6 μg/mL HDL at RT to attach to the cell surface. After 1 h, the cells were washed of free SARS-CoV-2pp, and then switched to 37 °C to allow SARS-CoV-2pp entry for another 3 h before harvesting for immunostaining. The scale bar indicates 10 μm. The data shown are representative of two independent experiments. c,d, 293T cells transfected with HA-SR-B1 were exposed to SARS-2-S in solutions containing 0.1% FBS or 6 μg/mL HDL at RT to attach to the cell surface. After 30 min, the cells were washed and detached before immunolabeling for flow cytometry (n=3). Percentage of SARS-2-S positive cells (c) or MFI of SARS-2-S (d) on the cell surface was quantified with isotype and SARS-2-S staining. Data are mean ± SEM. ***P < 0.001 by one-way ANOVA and Bonferroni’s post hoc analysis. e, Immunoprecipitation analysis in 293T cells transfected with Flag-SARS-2-S. Approximate molecular weight (kDa) marker positions are indicated to the left of the blot. Representative immunoblots are shown (n=2). f, Huh-7 cells preincubated with ITX 5061 (25 μM) or BLT-1 (25 μM) were inoculated with SARS-CoV-2pp in solutions containing 6 μg/mL HDL and pseudotyped viral entry was analyzed by luciferase activity at 48 h after infection (n=3). Signals obtained without compounds in 0.1% FBS were used for normalization. Data are mean ± SEM. ***P < 0.001 by one-way ANOVA and Bonferroni’s post hoc analysis. g, Huh-7 cells preincubated with ITX 5061 (25 μM) were infected with SARS-CoV-2 (MOI = 0.001) in solutions containing 6 μg/mL HDL at the indicated concentrations for 1 h and the cells were harvested to detect the gene copy number of virus with qRT-PCR at 24 h after infection (n=3). Data are mean ± SEM. ***P < 0.001 by two-tailed Student’s t-tests. h, Representative images of immunohistochemistry staining for ACE2 and SR-B1 was performed on the paraffin-embedded human normal organ tissue microarray with antibodies against ACE2 or SR-B1 and counterstained with hematoxylin to show nuclei (blue). Scale bars indicate 200 μm or 20 μm as indicated.

The implication of SR-B1 was further validated by testing the effects of known SR-B1 inhibitors on SARS-CoV-2 infection, including ITX 5061^13^ and block lipid transport-1 (BLT-1). BLT-1 is a small compound that inhibits the SR-B1-mediated selective transfer of lipids without affecting HDL-SR-B1 binding^14^. Both of these compounds exerted no unwanted cytotoxic effects (Extended Data Fig. 3a, b) and significantly blocked SARS-2-Spp entry (Extended Data Fig. 3c, d). Importantly, the HDL-mediated enhancement of SARS-CoV-2pp infectivity was significantly blocked by ITX 5061 and BLT-1 (Fig. 4f). To obtain direct evidence that clinical-grade ITX 5061 can indeed interfere with SARS-CoV-2 infections, we infected Vero E6 cells with SARS-CoV-2 in the presence of HDL with or without ITX 5061. Viral RNA was purified from cells and assayed by qRT-PCR as a marker for replication. Indeed, ITX 5061 treatment strongly inhibited SARS-CoV-2 infection stimulated by HDL (Fig. 4g). Therefore, SR-B1-mediated HDL uptake facilitates SARS-CoV-2 infection and posit potential therapeutic strategy.

Evidences of SARS-CoV-2 attacking multiple organs such as digestive, cardiovascular, urinary system and male reproductive systems have been reported, the distribution analysis of SR-B1 in the susceptible cells at the site of infection may reveal the viral tropism and potential transmission routes. Immunohistochemical studies showed cell surface staining of SR-BI at multiple regions of the human lung tissues, which coincides with the site of SARS-CoV-2 replication (Fig. 4h). Interestingly, the co-expression of SR-B1 and ACE2 could also be recognized in extrapulmonary tissues including colon, retina, small intestine, and testis (Fig. 4h and Extended Data Fig. 4a). Overall, our data demonstrated that SR-B1 was coexpressed with ACE2 on multiple susceptible tissues and could potentially be involved during SARS-CoV-2 infection.

## Discussion

With over 10 million infected people, COVID-19 represents a major public health problem of high socio-economic impact. However, treatment options for COVID-19 are limited, and a vaccine for prevention against COVID-19 infection is currently not available. The results of our present study demonstrated that SR-B1 is required for SARS-CoV-2 entry and infection. Indeed, SARS-CoV-2 entry is inhibited by silencing SR-B1 expression and by SR-B1 antagonists. Moreover, SR-B1 expression levels have been shown to affect SARS-2-Spp attachment, entry and authentic SARS-CoV-2 infection. Thus, as a key entry factor, SR-B1 offers a novel therapeutic strategy to prevent or cure SARS-CoV-2 infection.

HDL is one of the major groups of lipoproteins present in human plasma that nonspecifically blocks or facilitates viral entry through the SR-B1. In the present study, we observed that the presence of HDL significantly increases authentic SARS-CoV-2 infection. Mechanistically, HDL facilitates viral attachment and entry by associating with SARS-2-S. The NTD and RBD domains of SARS-2-S1 are involved in its interaction with HDL. An antibody blocking SARS-2-S1-HDL binding was observed to strongly inhibit HDL-enhanced SARS-2-Spp entry. To our surprise, we did not observe an interaction between SARS-2-RBD and cholesterol (data not shown). Thus, the key determinants of SARS-2-S1 in mediating HDL interaction and their implications in viral entry requires further investigation.

As a key entry cofactor, SR-B1 cannot bind to SARS-2-S protein directly. However, HDL-enhanced SARS-2-S attachment and SARS-CoV-2pp entry were greatly increased by SR-B1 expression. Importantly, SR-B1 expression conferred the greatest cell susceptibility to SARS-CoV-2pp only when coexpressed with ACE2. However, additional cell lines with varying degrees of SR-B1 and ACE2 receptor expression or primary targeted cells are needed to elucidate the function of SR-B1 in mediating viral entry. Our present data support the notion that SR-B1 functions as a linker molecule that recruits viral particles to come into close proximity with ACE2 by associating with HDL (Extended Data Fig. 5b). A major breakthrough will be the identification of the SR-B1-ACE2 membrane complex that is important for viral entry.

If SR-B1 is the primary receptor defining SARS-CoV-2 attachment to target cells, why does the lipid transfer activity of SR-B1 promotes viral entry? We know that SR-B1-mediated lipid transfer from HDL results in local increases in the cholesterol content of cell membranes^8,15^. Cholesterol and the abundant presence of cholesterol within lipid rafts play an essential role in viral entry and fusion^16-19^, and cholesterol depletion from cellular membranes has been shown to inhibit SARS-CoV-2 infection^20^. We thus speculated that SARS-CoV-2 may exploit SR-B1 physiological function to achieve its entry and fusion processes.

The widespread expression of multiple subsets of SR-B1 across diverse tissues may contribute to the high transmissibility of SARS-CoV-2. To understand how SR-BI and ACE2 coordinate viral entry, it is important to compare receptor expression pattern(s) and localization in lower respiratory tissues. In the current study, we demonstrated that lower respiratory tissues express SR-B1 and ACE2, consistent with their ability to support SARS-CoV-2 entry. Of note, although ectopic SR-B1 introduction did not modify ACE2 expression at 24 h after transfection, significantly increased intracellular and surface ACE2 protein levels were observed at 48 h after SR-B1 transfection (data not shown). It is thus interesting to uncover the spatiotemporal relationship between SR-B1, lipid raft, and ACE2. Publicly available patient cohorts indicated that expression of SR-B1 was greater in atherosclerotic arteries than normal arteries, we have reasons to believe that SR-B1 plays a major role in pathophysiology of COVID-19, especially in patients with severe acute metabolic disturbance.

ACE2 was identified as a functional SARS coronavirus receptor both in cell lines and in vivo^21^. Spike protein–mediated ACE2 downregulation then contributes to the severity of lung pathologies^22^. Commonalities between SARS-CoV-2 and SARS-CoV infection implicate the uses of antibodies, peptides and small compounds targeting ACE2 in the treatment of SARS-CoV-2. By identification of SR-B1 as a SARS-CoV-2 entry coreceptor, we expanded upon antiviral therapeutic approaches to include drugs that target SR-B1. Amazingly, ITX 5601, a clinically approved promising inhibitor of HCV infection, strongly inhibits SARS-CoV-2 infection in vitro. In summary, the results of the present study provide key insights into the remarkable capacity of the SARS-CoV-2 virus to hijack SR-B1 functions to promote its entry and defined potential antiviral approaches to interfere with viral entry.

## Methods

### Reagents and antibodies

His-tagged SARS-2-S (S1+S2 ECD; YP_009724390.1;Val16-Pro1213; 40589-V08B1), SARS-2-S1 (YP_009724390.1;Val16-Arg685; 40591-V08H), and SARS-2-S2 (ECD; YP_009724390.1; Ser686-Pro1213; 40591-V08B) proteins were from Sino Biological. ITX 5061(HY-19900) was from MedChemExpress. Cholesterol (C8667), campesterol (C5157), epicholesterol (R207349), BLT-1 (SML0059), HDL (L8039), DMEM (high glucose, D5796) and protease inhibitors (11206893001) were from Sigma-Aldrich. Human serum (A3969001) and FBS (A3160901) were from Gibco. Lipofectamine 2000 (11668-027) was from Invitrogen. A tissue Microassay (MNO661) was from Biomax. Monolith His-Tag Labeling kit (MO-L018) was from NanoTemper Technologies.

α-Tubulin for immunoblotting (T9026, clone DM1A; 1:1,000 dilution), HRP-labeled anti-Flag (A8592, clone M2;1:1,000 dilution), anti-human FITC conjugated-IgG (AP113F;1:50 dilution) and anti-HA (H9658, clone HA-7;1:20,000 dilution) were from Sigma-Aldrich. Antibodies against SR-B1 for immunoblotting (ab217318, clone EPR20190; 1:2,000 dilution), immunohistochemistry (ab217318,clone EPR20190; 1:100 dilution), and against ACE2 for immunoblotting (ab108252, clone EPR4435(2); 1:1,000 dilution) immunohistochemistry (ab108252, clone EPR4435(2); 1:50 dilution) were from Abcam. An antibody against ACE2 for FACS (21115-1-AP; 1:100 dilution) was from Proteintech. A PE anti-His antibody (362603, clone J095G46; 1;100 dilution) and an antibody against SR-B1 for FACS (363208, clone m1B9; dilution 1:100) were from Biolegend. The SARS-2-S mAb (GTX632604, clone 1A9; 1:1,000 dilution) was from GeneTex. The SARS-CoV-2 S1 antibody (40150-R007) for confocal microscopy was from Sino Biological. 1D2 mAb was from the laboratory of Wei Chen.

### Plasmids

Expression plasmids for human HA-SR-B1 (H3784), Flag-ACE2 (H3673) and HA-ApoA1 (H3912) were obtained from VigeneBiosciences. Expression plasmids for SARS-2-S, VSV-G and the HIV-1 NL4-3 ΔEnv Vpr Luciferase Reporter Vector (pNL4-3-Luc-R-E-) were obtained from the laboratory of Wei Chen. The codon-optimized (for human cells) SARS-2-S gene was based on Genebank ID: QHD43416.1. All constructs were sequence verified. The siRNAs targeting human genes were obtained from Invitrogen and had the following target sequences:

siSR-B1-1: 5’-CAAGUUCGGAUUAUUUGCUTT-3’

siSR-B1-2: 5’-CAUGAUCAAUGGAACUUCUTT-3’

siSR-B1-3: 5’-GCCUCUACAUGAAAUCUGUTT-3’

Scrambled siRNA oligonucleotides from siSR-B1-3 were used as a control (siCtrl).

Cell culture and transfection

293T (CRL-3216), Vero E6 (CRL-1568), MDCK (CCL-34) and Hepa 1-6 (CRL-1830) cell lines were from the American Type Culture Collection (ATCC, Rockville, MD, USA). Huh-7 (0403) was from Japanese Collection of Research Bioresources. HeLa-hACE2 (012) was from Beijing Vitalstar Biotehnology. All cell lines were previously tested for mycoplasma contamination and incubated in Dulbecco’s modified Eagle’s medium at 37 °C under a humidified atmosphere with 5% CO_2_. All media were supplemented with 10% FBS, 100 U/mL of penicillin, 0.1 mg/mL of streptomycin, 1× nonessential amino acid solution and 10 mM sodium pyruvate. Lipofectamine 2000 was used for transfection following the manufacturer’s protocol.

### Microscale thermophoresis (MST)

MST assays were conducted as previously described^23^. SARS-2-S-His-tagged, SARS-2-S1-His-tagged, and SARS-2-S2-His-tagged proteins were labeled with NT-647 dye for 30 min at room temperature, as recommended by the Monolith His-Tag Labeling Kit RED-tris-NTA protocol (NanoTemper Technologies, MO-L008). The binding buffer used for reactions [50 mM Tris-HCl (pH 7.5), 150 mM NaCl, and 1 mM DTT] contained micelles of 0.5% fos-choline 13, and a 16-step, 2-fold dilution curve of cholesterol and cholesterol derivatives ranged from 4 μM to 122 pM. Cholesterol or other sterol dry powders were mixed with binding buffer at 25 °C with mild agitation for 3 h. Labeled protein (100 nM, final concentration) in binding buffer was then added to a dilution curve of cholesterol or cholesterol derivatives at room temperature. Samples were loaded into standard glass capillaries (NanoTemper Technologies, MO-K022). Microscale thermophoresis was completed in three independent experiments on a Monolith NT. 115 instrument (NanoTemper Technologies) running MO.Control software with settings of 60-80% excitation power and 40% MST power at room temperature.

In the HDL-binding assay, labeled protein (100 nM, final concentration) in binding buffer (145 mM NaCl) was added to a dilution curve of HDL at room temperature, where the dilution curve of HDL ranged from 6.25 μg/mL to 190.7 pg/mL (16-step, 2-fold). Samples were loaded into standard glass capillaries (NanoTemper Technologies, MO-K022). Microscale thermophoresis was completed in 3 independent repeat experiments on a Monolith NT.115 instrument (NanoTemper Technologies), running MO.Control software with settings of 60-80% excitation power and 40% MST power at room temperature.

In the competition assay, 100 nM of labeled SARS-2-S in binding buffer was incubated with 400 nM cholesterol either alone or with unlabeled SARS-2-S at the indicated concentrations.

Original data were obtained from MO.Affinity Analysis software. Fluorescence intensity values were averaged and expressed as the relative change over the results from the 0 μM ligand condition. The data were to fitted saturation binding equations using GraphPad Prism 8.0. Dotted lines indicate where data could not be fitted.

### Peptide-cholesterol binding assays

The following synthetic peptides, encompassing the CARC-CRAC region of SARS-2-S, were purchased from Scilight-Peptide. A control peptide with random amino acid sequence was also included in the assay:

Control (Ctrl): KKKARVRIFLYGFLLQLLMPVWTMKKK

24-42: KKKLPPAYTNSFTRGVYYPDKVKKK

129-150: KKKKVCEFQFCNDPFLGVYYHKNNKKKK

267-277: KKKVGYLQPRTFLLKKK

445-461: KKKVGGNYNYLYRLFRKSNLKKK

776-806: KKKKNTQEVFAQVKQIYKTPPIKDFGGFNFSQILKKK

1203-1218: KKKLGKYEQYIKWPWYIWLKKK

The N-term and C-term Lys residues increase the solubility of the hydrophobic CARC-CRAC transmembrane segment. Replacing the Pro residue located between the CARC and the CRAC with Trp conferred intrinsic fluorescence upon 280 nm excitation, with maximum emission at 350 nm. As previously reported, the conformation of the peptide affects the 350 nm emission intensity, and the fluorescence titration assay was performed as previously described (6). Stock solutions of 5 mM peptide in DMSO were diluted to a final concentration of 5 μM in assay buffer containing 140 mM NaCl and 20 mM Tris (pH 8.5) supplemented with 2 mM TCEP and 2% (v/v) EtOH. Stock solutions of cholesterol in EtOH were diluted to final concentrations ranging between 0.5 and 50 μM in 500-μL volumes of peptide solution, which were incubated overnight at 25 °C with mild agitation. Fluorescence measurements were performed in triplicate on a Cary Eclipse fluorescence spectrophotometer (Agilent Technologies), with excitation and emission performed at 280 and 350 nm, respectively. All measured fluorescence values fell within the linear range of the instrument. Fluorescence intensity values were averaged and expressed as relative change over the 0 μM ligand condition. Data were fitted to saturation binding equations using GraphPad Prism 6. Dotted lines indicate where data could not be fitted.

### Pseudovirus production

All pseudoparticles were generated in 293T cells transfected with the HIV backbone vector pNL4-3.Luc.R-E together with the expression plasmids SARS-2-S or VSV-G. Pseudoparticle-containing supernatants were harvested at 48 h and frozen to −80 °C after centrifugation at 2000×g for 5 min.

### Pseudovirus entry assays

Cells were cultured in 96-well transparent-bottom flat culture plates (#3603, Corning) precoated with poly-D-lysine. To assess the inhibitory activity of chemicals, the indicated concentrations of chemicals were added to cells 4 h prior to infection. To determine the neutralization ability of antibodies, serially diluted antibodies were added to cells 1 h prior to infection. SARS-2-Spp preincubated with HDL at 25 °C for 2 h were applied to the cells pretreated with chemicals or antibodies and infected at 37 °C for 24 h. After SARS-2-Spp infection, the efficiency of viral entry was determined through a firefly luciferase assay. Specifically, cells were washed with PBS once, and then 16 μL of Cell Culture Lysis Reagent (E153A, Promega) was added to each well. Then, the plate was incubated for 15 min with rocking at RT, after which cell lysate (8 μL) from each well was added to a 384-well plate (#3574, Corning) followed by the addition of 16 μL of Luciferase Assay Substrate (#E151A, Promega). Luciferase activity measurements were performed on a Spark 20M multimode microplate reader (Tecan). All infection experiments were performed in a biosafety level-2 (BSL-2) laboratory. Cells without viruses, HDL and chemicals were used as blank controls, and cells without chemicals and HDL were used as viral controls. All the infection experiments were performed in a biosafety level-2 (BSL-2) laboratory.

### Western blot analysis and immunoprecipitation

Cells were lysed in NP40 cell lysis buffer with fresh protease inhibitors. Supernatants were separated by SDS-PAGE gel after centrifugation and transferred to PVDF membranes for immunoblotting analyses using the indicated primary antibodies. Then, an aliquot of the total lysate (5%, v/v) was included as a control for the interaction assay, and immunoprecipitation was performed with the indicated antibodies. Then, anti-rabbit or anti-mouse HRP-conjugated antibodies were applied after 3 washes and the antigen-antibody complexes were visualized using chemiluminescence.

### Infection with authentic SARS-CoV-2

The SARS-CoV-2 strain (2019-nCoV BetaCoV/Beijing/AMMS01/2020) used in the present study was isolated from the lung lavage fluid of an infected patient and preserved in the State Key Laboratory of Pathogen and Biosecurity at Beijing Institute of Microbiology and Epidemiology. All the experiments involving viruses were performed in the BSL-3 Laboratory of the Beijing Institute of Microbiology and Epidemiology.

For HDL: Vero cells (1.6 ×10^5^ cells/mL) were infected with the SARS-CoV-2 isolate (MOI=0.001) preincubated with the indicated concentrations of HDL at 37 °C for 1 h. For chemicals and antibodies, Vero E6 or Huh-7 cells (1.6 ×10^5^ cells/mL) were cultured in a 96-well plate at 37 °C overnight. Then, the supernatant was discarded, and 100 μL of medium containing different concentrations of drugs was added to the wells of the plates and were incubated for 4 h (chemicals) or 1 h (for antibodies) prior to infection. Then, the cells were infected with the SARS-CoV-2 isolate (MOI=0.001) preincubated with the indicated concentration of HDL at 37 °C for 1 h.

For SR-B1 expression, Vero cells (1.6 ×10^5^ cells/mL) were transfected with a plasmid encoding SR-B1, while Huh-7 cells (1.6 ×10^5^ cells/mL) were transfected with siRNA oligos for SR-B1. After 24 h, cells were washed with PBS and infected with the SARS-CoV-2 isolate (MOI=0.001) that was preincubated with the indicated concentration of HDL at 37 °C for 1 h.

After 1 h of infection, the above cells were washed three times with PBS before 200 μL of DMEM medium was added. Then, at 24 h post infection, cells were harvested for viral RNA extraction with a viral RNA kit (52906, QIAGEN) according to the manufacturer’s instructions and detected by RT-qPCR assays with a One-Step PrimeScript RT-PCR kit (Takara, Japan) using SARS-CoV-2-specific primers on an Applied Biosystems 7500 Real-time PCR System.

The TaqMan primers for SARS-CoV-2 were as follows: 5’-TCCTGGTGATTCTTCTTCAGG-3’ and 5’-TCTGAGAGAGGGTCAAGTGC-3’. The SARS-CoV-2 probe used was as follows: 5’-FAM-AGCTGCAGCACCAGCTGTCCA-BHQ 1-3’.

Copies per milliliter were determined using a synthetic RNA fragment to amplify the target region.

### Flow cytometry for surface SR-B1 or ACE2 expression analysis

Cells were washed with PBS 2 times and incubated with APC-conjugated (APC anti-SR-B1) antibody (1:100) or anti-ACE2 antibody (1:100) followed by a secondary incubation with donkey-anti-rabbit antibody (1:200) conjugated to Alexa Flour 488. Measurement of cellular fluorescence was determined by FACS. Thirty thousand events were registered for each experiment. Cellular fluorescence was quantitated by mean fluorescence intensity (MFI) or percent positive cells.

### Binding assay

For SARS-2-S-cell surface binding, cells or cells transfected with HA-SR-B1 were detached in PBS and washed with PBS at 24 h after transfection. SARS-2-S proteins (140 nM) and HDL (6 μg/mL) preincubated at 37 °C for 1 h were then added to cells. After incubating at RT for 30 min, the cells were washed two times with PBS containing 2% FBS and then incubated for 30 min at RT with a PE-conjugated antibody (PE anti-His tag). Then, the cells were washed again with PBS containing 2% FBS, after which binding was detected by flow cytometry. The cells were analyzed on an Invitrogen Attune NxT Acoustic Focusing cytometer using Flowjo V10.0.7. At least 10,000 cells were analyzed for each sample.

### Internalization assay

To monitor SARS-2-Spp entry, 293T cells transfected with GFP-SR-B1 were detached in PBS and washed with PBS at 24 h after transfection. SARS-2-Spp preincubated with HDL at 37 °C for 1 h was applied to the cells at RT for 1 h and then shifted to 37 °C for 3 h. The cells were then washed with PBS two times, after which they were fixed, permeabilized, and blocked with 3% BSA. SARS-CoV-2 (2019-nCoV) Spike S1 antibody, rabbit MAb and Alexa Fluor 555-labeled donkey anti-rabbit IgG (H+L) were used to bind SARS-2-Spp. Virion localization was assayed by immunofluorescence microscopy.

### Immunohistochemistry

Immunohistochemical staining for ACE2 and SR-B1 was performed on the paraffin-embedded human normal organ tissue microarray (MNO661, Biomax). Antigen retrieval was performed by placing the slides in 10 mM sodium citrate buffer (pH 6.0) and maintained at a subboiling temperature for 10 min. After blocking in 1% normal goat serum, the sections were incubated with ACE2 or SR-B1 monoclonal antibodies at 4□°C overnight in a humidified chamber, which was followed by an incubation with HRP-labeled goat anti-mouse IgG secondary antibody (Beijing ZSGB Biotechnology, ZDR-5307). Subsequently, the sections were incubated with a goat anti-rabbit IgG secondary antibody (HRP) (Beijing ZSGB Biotechnology, PV9001) for 60 min and then visualized with 3,30-diaminobenzidine tetrahydrochloride (DAB). The slices were viewed under an Olympus microscope.

### Statistical methods

In the present study, GraphPad Prism 8.0 was used for statistical calculations and data plotting. Differences between two independent samples were evaluated by two-tailed Student’s t-tests. Differences between multiple samples were analyzed by one-way ANOVA followed by Bonferroni’s post hoc analysis. All tests were two-tailed, unless otherwise indicated. We considered a *P* value ≤ 0.05 to be statistically significant. Significance values were set as follows: ns (not significant), P > 0.05; *, P < 0.05; **, P < 0.01; ***, P < 0.001.

### Data availability

The data that support the findings of this study are available from the corresponding author upon reasonable request. Source data for Figs. 1-4 and Extended Data Figs. 1-4 are presented with the paper.

## Acknowledgements

The authors would like to thank the National Key Research and Development Program of China (2018YFA0900800), National Natural Science Foundation of China (81671973, 31670761, 31872715, and 81773205), Beijing Natural Science Foundation (5182029, 5182030), Shandong Provincial Natural Science Foundation of China (No.ZR2015HM003) and Natural Science Foundation of China (No.81572620) for the support. The authors would like to thank Dr. Qing Chang for MST data collection at the Protein Preparation and Characterization Platform of the Tsinghua University Technology Center for Protein Research.

## Author contributions

CW. W., LM. W., QL. Y., XL. W., J. Z., YH. Z., J. S., XP. Y., J. G., H. Y., HL. L., WC. Z., YM. P. and HT. L. performed the experiments. C. F., XL. Y., YF. W., XJ. W., JM. L., Z. Z., R. W., P. D., YL. Z., F. Y., provided reagents and intellectual input. CW. W., R.Z., W. C., Q. G., Y. C. and H. Z. designed and coordinated the study and wrote the manuscript. All authors read and approved the manuscript.

