## supplemental material for "SARS-CoV-2 manipulates the SR-B1-mediated HDL uptake pathway for its entry"

**Supplementary Figures and Figure Legends**

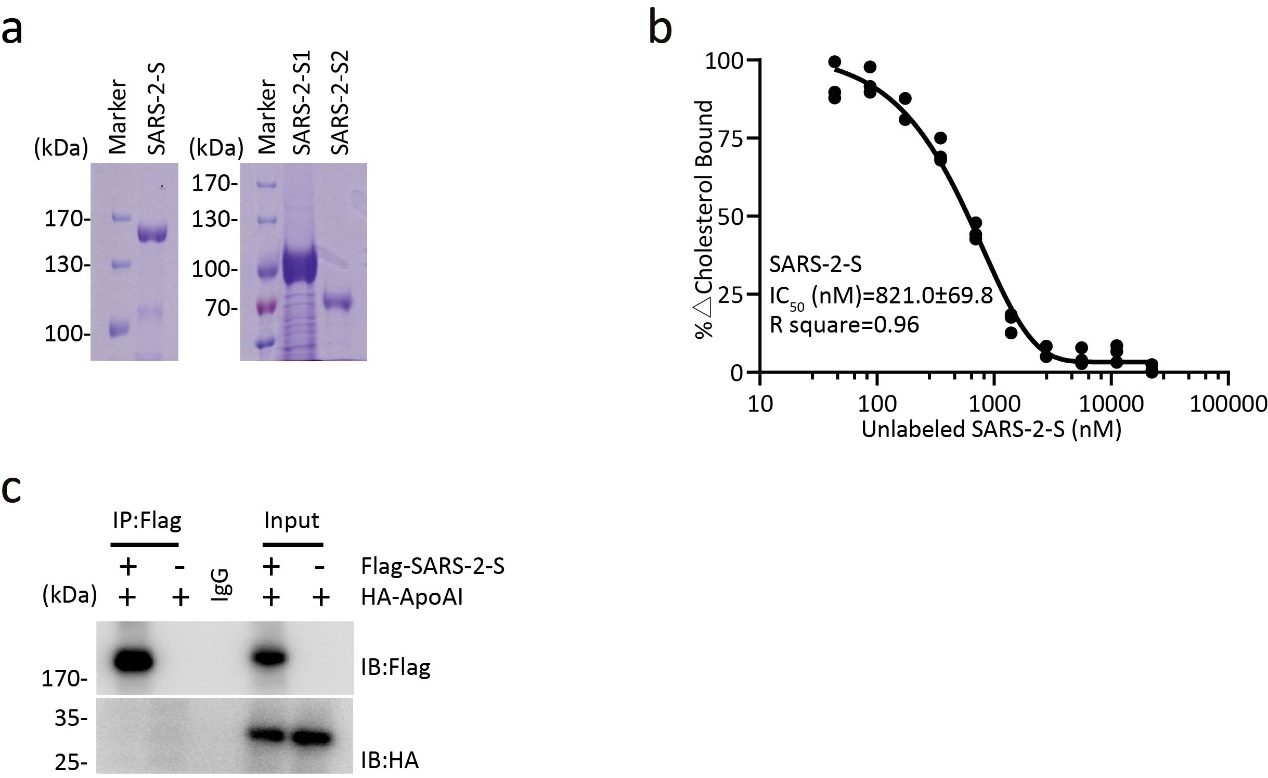

**Extended Data Fig. 1 The spike protein of SARS-CoV-2 binds to cholesterol and HDL.** a, Validation of SARS-2-S, SARS-2-S1 and SARS-2-S2 protein by SDS-PAGE. b, Competitive binding of the indicated concentrations of unlabeled SARS-2-S with NT-647-NHS dye-labeled SARS-2-S in 0.5% fos-choline 13 micelles containing 200 nM cholesterol (n=3). IC_50_ was determined by Hill slope. c, Immunoprecipitation analysis in 293T cells transfected with Flag-SARS-2-S and HA-ApoA1. Approximate molecular weight (kDa) marker positions are indicated to the left of the blot. Representative immunoblots are shown (n=3).

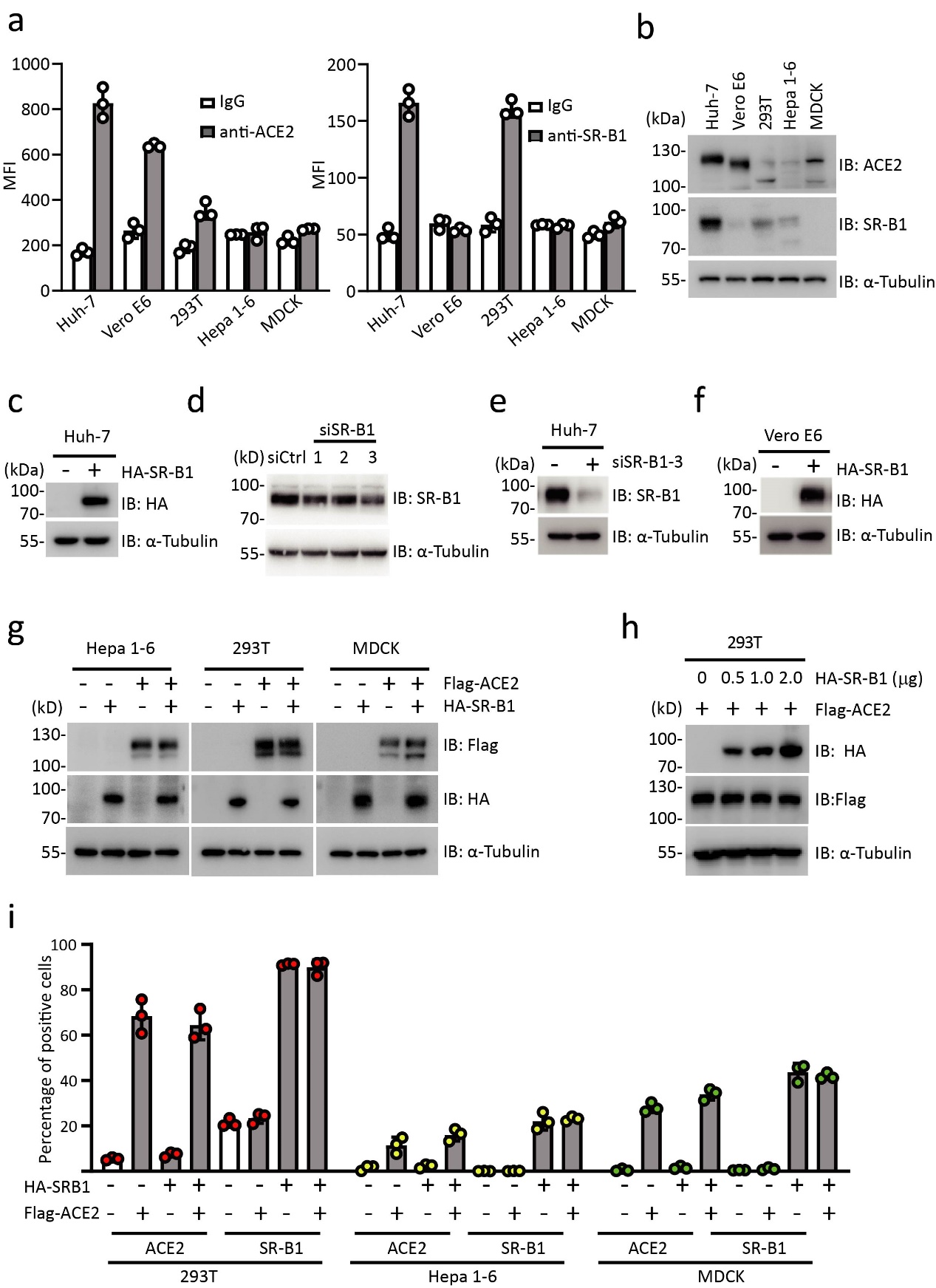

**Extended Data Fig. 2 SR-B1 expression confers susceptibility to SARS-CoV-2 infection.** a,b, Flow cytometry (a) or immunoblotting (b) analysis of ACE2 and SR-B1 expression in the indicated cell lines. c,d, Immunoblotting analysis of SR-B1 expression in Huh-7 cells transfected with HA-SR-B1 from Fig. 3a (c) or transfected with three siRNA oligos for SR-B1 from Fig. 3b (d). α-Tubulin was used as the equal loading control. No. 3 oligos (siSR-B1-3) were used for the pseudotyped viral infection assay. e,f, Immunoblotting analysis of SR-B1 expression in Huh-7 cells transfected with siSR-B1-3 from Fig. 3c (e) or Vero E6 cells transfected with HA-SR-B1 from Fig. 3d (f). α-Tubulin was used as the equal loading control. g,h, Immunoblotting analysis (g) or flow cytometry analysis (h) ACE2 and SR-B1 expression in the indicated cell lines transfected with plasmids encoding ACE2 and or SR-B1 from Fig. 3e, f, g. i, FACS analysis of ACE2 and SR-B1 expression in 293T cells transfected with Flag-ACE2 together with increasing concentrations of HA-SR-B1 from Fig. 3h.

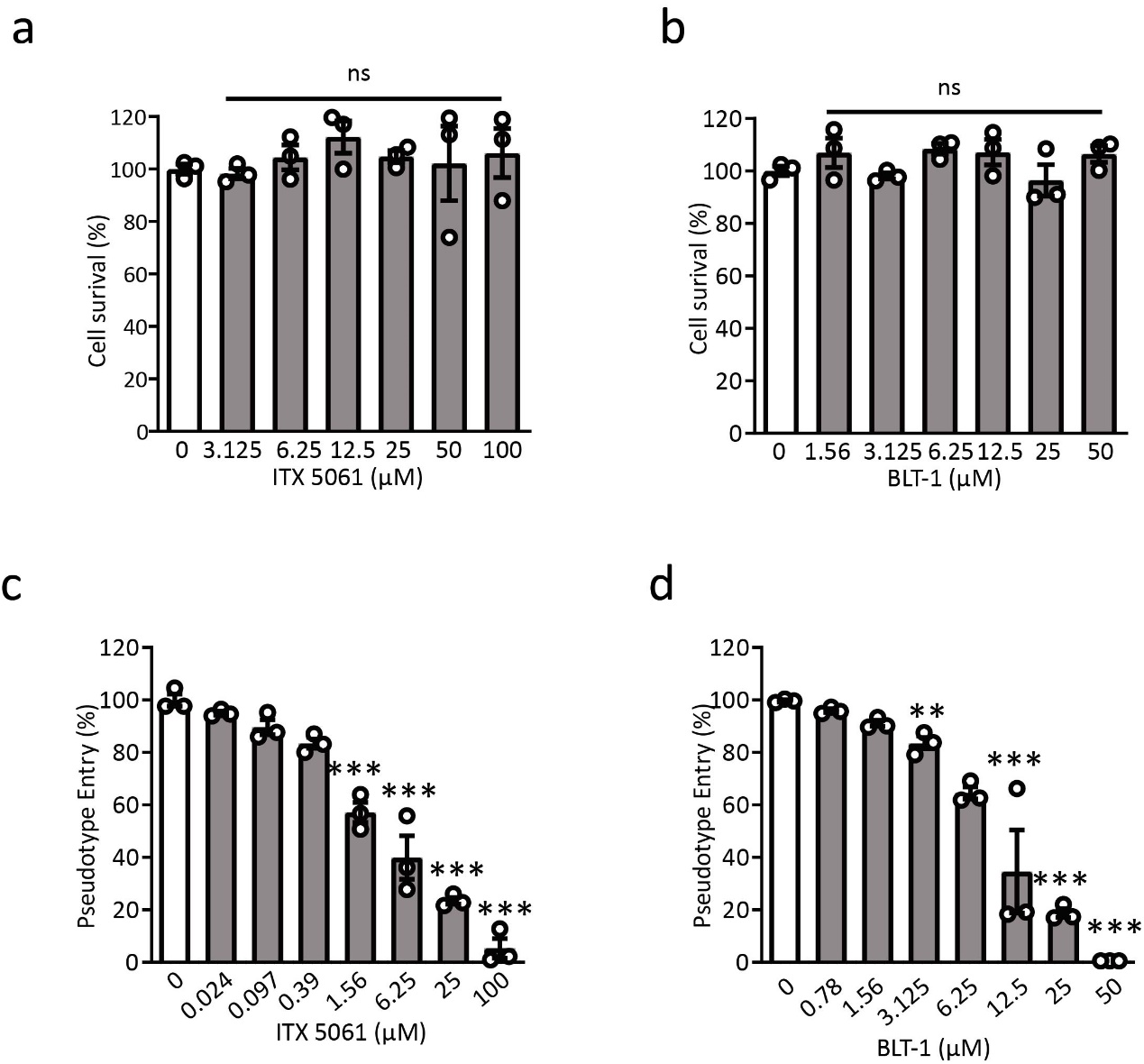

**Extended Data Fig. 3 SR-B1 antagonists block SARS-2-Spp infection.** a,b, Huh-7 cells preincubated with the indicated concentrations of ITX 5061 (a) or BLT-1 (b) were inoculated with SARS-CoV-2pp in 10% FBS and pseudotyped viral entry was analyzed by luciferase activity at 48 h after infection (n=3). Signals obtained without compounds were used for normalization. Data are mean ± SEM. **P < 0.01, ***P < 0.001 by one-way ANOVA and Bonferroni’s post hoc analysis. c,d, Huh-7 cells preincubated with the indicated concentrations of ITX 5061 (c) or BLT-1 (d) for 48 h and MTT was added to the cells to measure OD_492_ value (n=3). Signals obtained without compounds were used for normalization. Data are mean ± SEM. ns by one-way ANOVA and Bonferroni’s post hoc analysis.

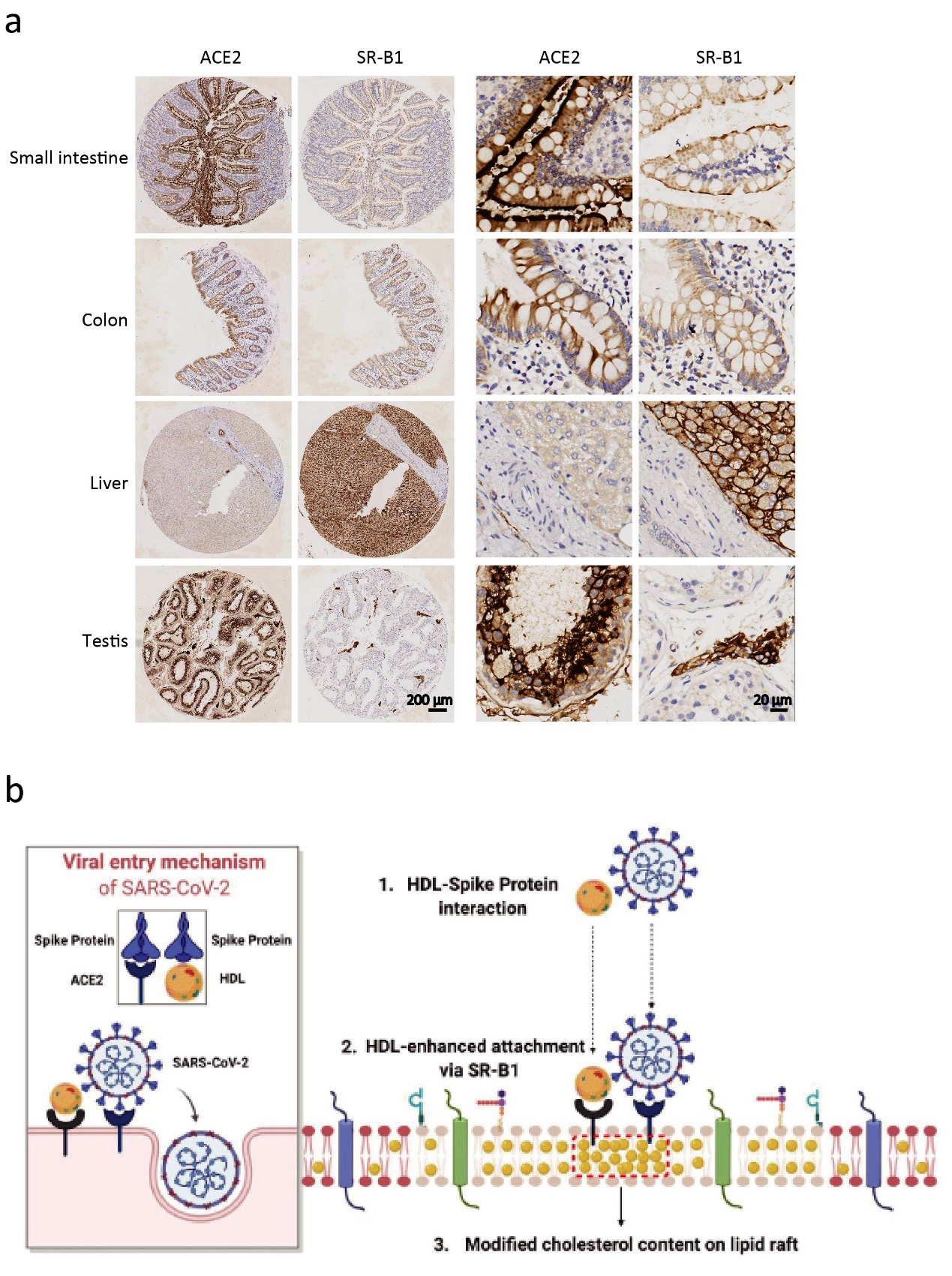

**Extended Data Fig. 4 Coexpression of SR-B1 and ACE2 in multiple normal human tissues.** a, Representative images of immunohistochemistry staining for ACE2 and SR-B1 was performed on the paraffin-embedded human normal organ tissue microarray with antibodies against ACE2 or SR-B1 and counterstained with hematoxylin to show nuclei (blue). Scale bars indicate 200 μm or 20 μm as indicated. b, Hypothetical model of SR-B1 involved in the entry of SARS-CoV-2. The figure was created and exported with BioRender.com under a paid subscription.

A figure exemplifying the gating strategy is provided.

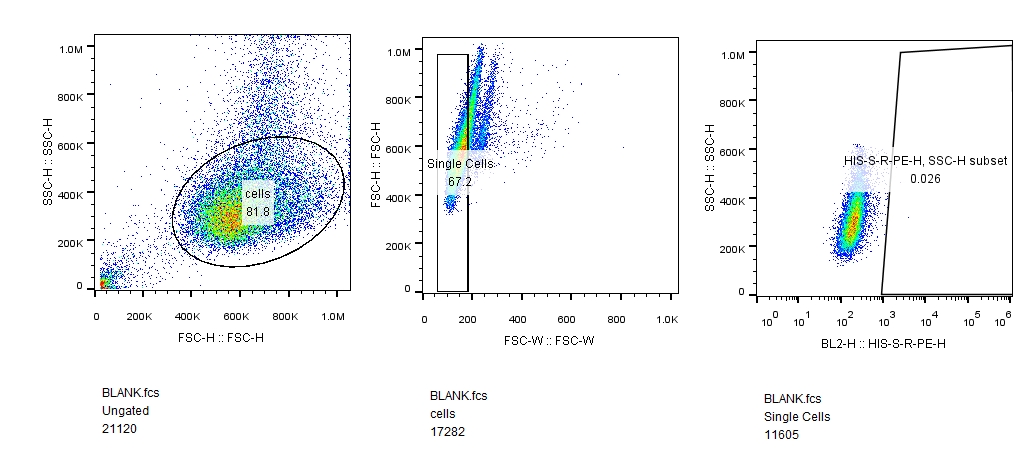
